# RapTCR: Rapid exploration and visualization of T-cell receptor repertoires

**DOI:** 10.1101/2023.09.13.557604

**Authors:** Vincent M.L. Van Deuren, Sebastiaan Valkiers, Kris Laukens, Pieter Meysman

**Affiliations:** Adrem Data Lab, Department of Computer Science, University of Antwerp, Antwerp, Belgium; AUDACIS, Antwerp Unit for Data Analysis and Computation in Immunology and Sequencing, University of Antwerp, Antwerp, Belgium

**Keywords:** T-cell receptor (TCR), immune repertoire visualization

## Abstract

**Motivation:** The acquisition of T-cell receptor (TCR) repertoire sequence data has become faster and cheaper due to advancements in high-throughput sequencing. However, fully exploiting the diagnostic and clinical potential within these TCR repertoires requires a thorough understanding of the inherent repertoire structure. Hence, visualizing the full space of TCR sequences could be a key step towards enabling exploratory analysis of TCR repertoire, driving their enhanced interrogation. Nonetheless, current methods remain limited to rough profiling of TCR V and J gene distributions. Addressing this need, we developed RapTCR, a tool for rapid visualization and post-analysis of TCR repertoires.

**Approach:** To overcome computational complexity, RapTCR introduces a novel, simple embedding strategy that represents TCR amino acid sequences as short vectors while retaining their pairwise alignment similarity. RapTCR then applies efficient algorithms for indexing these vectors and constructing their nearest neighbor network. It provides multiple visualization options to map and interactively explore a TCR network as a two-dimensional representation. Benchmarking analyses using epitope-annotated datasets demonstrate that these RapTCR visualizations capture TCR similarity features on a global level (e.g., J gene) and locally (e.g., epitope reactivity). RapTCR is available as a Python package, implementing the intuitive scikit-learn syntax to easily generate insightful, publication-ready figures for TCR repertoires of any size.

**Availability and Implementation:** RapTCR was written in Python 3. It is available as an anaconda package (https://anaconda.org/vincentvandeuren/raptcr), and on github (https://github.com/vincentvandeuren/RapTCR). Documentation and example notebooks are available at vincentvandeuren.github.io/rapTCR_docs/.

## 1 Introduction

Our adaptive immune system fights hypervariability with hypervariability: to recognize and respond to the plethora of continually mutating pathogens, it relies on an extremely diverse set of immune receptors. In T-cells, the key molecule orchestrating T-cell reactivity and activation are their T-cell receptors (TCRs), which are uniquely generated during the V(D)J recombination process. TCRs continually interact with peptides presented by major histocompatibility complexes (pMHCs). This intricate interplay between TCRs and pMHCs forms the basis of antigen recognition and enables the immune system to mount highly specific and effective immune responses against a wide range of pathogens. Hence, monitoring the collection of all TCRs within an individual – the TCR repertoire – can provide unique insights into adaptive immunity and the antigen exposures that shape this repertoire. Recent advances in single cell and high-throughput immunosequencing methods have provided the means to characterize the TCR repertoire in unprecedented detail (Robins et al., 2009; Rosati et al., 2017). Currently, one of the major challenges is the subsequent post-analysis of these immune sequencing data, which mostly remains hindered by the adaptive immune systems’ inherent nature – its immense diversity.

To better understand T-cell receptor (TCR) repertoires, it’s crucial to recognize their inherent structure. This insight hinges on the fact that TCRs sharing similar sequences or motifs likely target the same epitopes, as supported by various studies (Glanville et al., 2017; Dash et al., 2017; Meysman et al., 2018). Consequently, we can envision the TCR repertoire as a network, where each TCR serves as a node and edges linking highly similar TCRs (Madi et al., 2017; Priel et al., 2018). Such an approach has demonstrated its effectiveness in studies that cluster highly similar sequences in the context of infection with SARS-CoV-2, hepatitis B, or tuberculosis (Mayer-Blackwell et al., 2021; Elias et al., 2022; Musvosvi et al., 2023). Visualizing the topology of these TCR repertoire networks holds the potential to provide deeper insights into their underlying characteristics, including epitope reactivity. However, a suitable method for visualizing TCR repertoire networks and exploring immune repertoires remains elusive. Current repertoire-level visualization methods are limited to plotting V- and J-segment usage or CDR3 length distributions (Dash et al., 2017; Ni et al., 2020; ImmunoMind Team, 2019; Nazarov et al., 2022), neglecting the intricacies of TCR diversity. Specialized TCR distance metrics have been introduced (Dash et al., 2017; Chronister et al., 2021); however, their use becomes impractical for large repertoires, as their computational complexity scales quadratically with repertoire size. Methods that tackle these computational challenges, e.g. TCR clustering techniques such as ClusTCR (Valkiers et al., 2021) are ill-suited to serve as the foundation for repertoire visualization, as they focus on highly similar, same-length CDR3 sequences.

With RapTCR, we created an easy-to-use Python toolkit that overcomes these hurdles and enables visualization of TCR repertoires. At the core of RapTCR is a novel and straightforward embedding method, which transforms TCR sequences into fixed-length vectors while preserving their pairwise similarities. This efficient representation enables optimized strategies for finding nearest neighbors and reducing dimensionality, thereby enabling visualization and interrogation of TCR repertoires of any size. Importantly, our work demonstrates that these visualizations convey meaningful insights, capturing both global and local features of the TCR sequence space. RapTCR is available as a Python package, empowering researchers to effortlessly generate insightful, publication-ready figures for TCR repertoires of any scale.

## 2 Approach

RapTCR can read in processed TCR sequencing data from various popular formats, including TSV files that comply to the AIRR standards (Heiden et al., 2018). Data is read as a Repertoire object, and further analysis is enabled by a user-friendly, scikit-learn-like syntax.

A widely used method to measure the similarity between sequences is Needleman-Wunsch alignment with a BLOSUM62-based scoring. However, when dealing with large datasets, scoring the similarity between every pair of sequences becomes computationally infeasible. Instead, RapTCR employs a straightforward strategy to represent TCR sequences as a vector of fixed length, with the Euclidean distance between TCR vectors approximating the alignment-based score. The embedding vector is the result of a multiplication of two matrices, which respectively encode information on the TCR amino acid composition and similarity, and the position of thee amino acids in the sequence (Figure S1). Our embedding approach is designed so that closely related sequences (with a high pairwise alignment score) also have a low Euclidean distance in the embedding space (Figure S2). Hence, these embedded TCR CDR3 vectors enable RapTCR to efficiently constuct the approximate k-nearest neighbor (k-NN) graph by employing highly optimized algorithms, such as NNdescent (Dong et al., 2011) and Faiss (Johnson et al., 2017). To further increase precision, more demanding TCR distance metrics can then be computed only for these neighboring pairs, pruning edges above an adjustable threshold distance. In this way, RapTCR drastically limits the number of exact pairwise distance computations needed, allowing it to construct the TCR similarity network at linear speed and with near-perfect recall (Figure S3).

For small repertoires, directly visualizing this graph representation can already be very informative. Nevertheless, RapTCR also offers several dimensionality reduction methods that enable the visualization of datasets of any size, facilitating their inspection, exploration, and interpretation. As a first option, RapTCR can visualize the repertoire graph as a minimum spanning tree (MST), by applying Kruskal’s algorithm and a force-directed drawing layout as described in Probst and Reymond (2020). Alternatively, the k-NN graph can be summarized using the uniform manifold approximation and projection (UMAP) algorithm (McInnes et al., 2018), for which we additionally provide a fast, pre-trained parametric UMAP model. These plots provide an overview of the entire CDR3 space, and preserve both global and local similarity features (Figure S4).

Lastly, RapTCR aims to be adaptable, seamlessly integrating with frequently-used immunoinformatic tools. It can handle paired-chain data (e.g. from a scanpy (Wolf et al., 2018) object), and clustered TCR sequence data (e.g. from ClusTCR (Valkiers et al., 2021)). The latter approach allows users to condense sequences to the cluster-level, enabling visualization of any repertoire, unrestricted by size. For all visualization methods, RapTCR effortlessly creates customizable, publication-ready figures, where different layers of information and annotations can be easily overlaid. For each of these plots, we additionally offer a web-based visualization option based on the plotly.js framework, where users can interactively explore and gain insight into their data.

**Figure 1:**
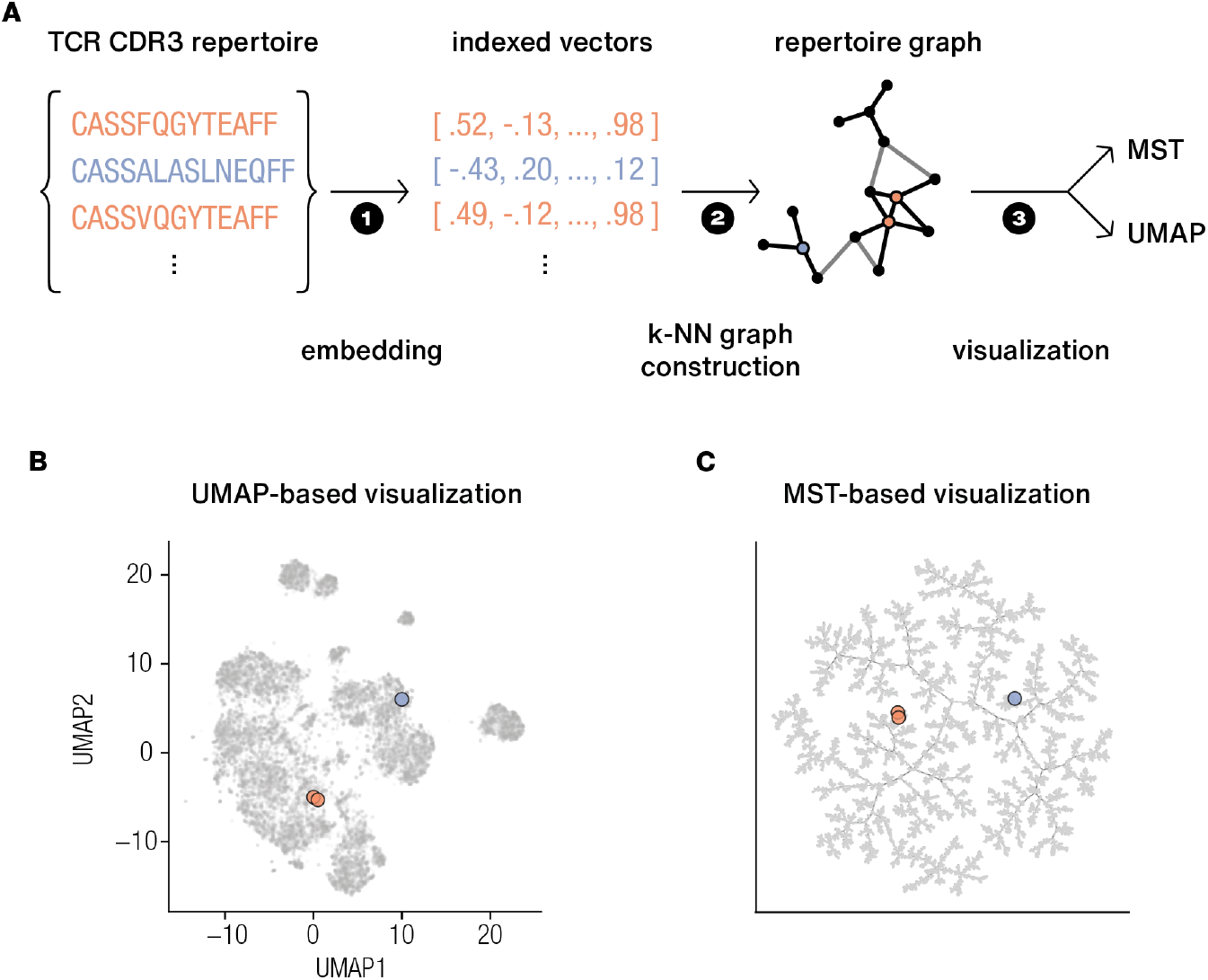
Overview of the general RapTCR workflow. **A**. The RapTCR methodology consists of three steps: (1) creating a vector embedding of each TCR sequence, which it uses to efficiently (2) find the nearest neighbors for each TCR, hence constructing the k-nearest neighbor graph, and (3) visualizing this graph using various dimensionality reduction techniques, depending on the dataset size. **B, C**. Example output of the UMAP-based visualization, and the MST-based visualization, respectively. Similar TCR sequences, such as the pair highlighted in orange, are located close together in the vector space, repertoire graph, and ultimately in the visualized TCR space.

## 3 Conclusion

Here, we introduced RapTCR, a Python toolkit designed to overcome the challenges of visualizing TCR repertoires, providing researchers with an accessible and efficient solution for exploring and interpreting these complex datasets. RapTCR builds upon an efficient way to find highly similar TCRs as defined by a global alignment-based metric. Other, more specialized TCR distance metrics (e.g. TCRdist Mayer-Blackwell et al. (2021)) have also been developed, and we aim to also provide an adapted embedding approach optimized for such metrics in future releases. We show that important TCR features, e.g. V/J gene information and epitope reactivity remain preserved in this low dimensional space. Nevertheless, we plan to add further informative layers of information to aid the interpretation of the resulting TCR repertoire map, e.g. highlighting the regions where it is enriched or depleted. In this way, we envision RapTCR as a key tool for exploratory TCR repertoire research, empowering researchers to unlock the wealth of information contained within these complex datasets and gain a deeper understanding of adaptive immunity.

## 4 Acknowledgements

This work was supported by the Research Foundation Flanders [1S40321N to Sebastiaan Valkiers].

## 5 Supplementary figures

**Figure S1:**
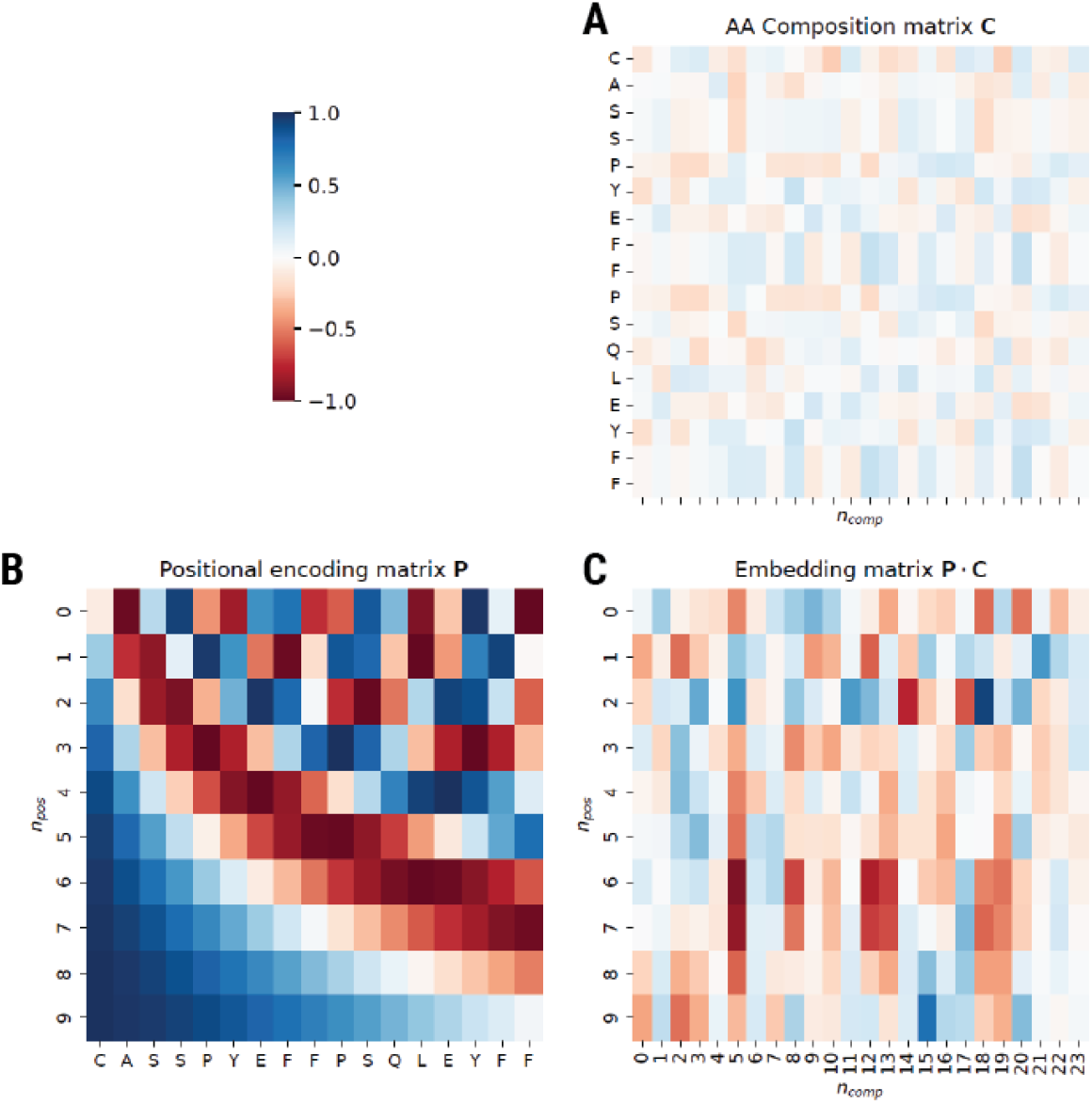
TCR sequence embedding. To embed a given TCR sequence as a fixed-length vector, we compute the product of two matrices, containing complimentary information. The amino acid composition matrix (**C**) encodes features of amino acid similarity, while the second matrix **P** adds information on the position of these amino acids in the TCR sequence. **A. Positional encoding (P)** For a TCR sequence of length *l* and with a specified number of positional dimensions *n*_pos_, we compute the positional encoding matrix **P** as the outer product of two vectors, **u** and **v**. The vector **u** consists of a geometrically decreasing sequence of length *n*_pos_. Each element is calculated as 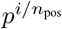, where *i* ranges from 0 to *n*_pos_ *−*1, and *p* is a user-defined parameter (default value: 0.4). The second vector **v** is a linear sequence of length *l*, ranging from 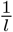 to 1. The positional encoding matrix is then computed as **P** = cos(2*rπ ·* **u** *⊗* **v**), where *r* is another user defined parameter (default value: 4.5). **B. Amino Acid Composition (C)** To construct the amino acid composition matrix **C**, we represent amino acids using a vector encoding derived from the BLOSUM62 substitution matrix (Henikoff and Henikoff, 1992). The conversion from BLOSUM62 log odds ratios to amino acid distances (*D*_*ab*_) is achieved using the equation 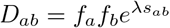, where *f*_*a*_ and *f*_*b*_ are the background frequencies for amino acids *a* and *b*, and *λ* is set to 0.347. Next, we apply a multidimensional scaling (MDS) function *f* to the resulting distance matrix *D* to create a *n*_comp_-dimensional projection for each amino acid. These projections preserve the amino acid similarity information from *D* in their pairwise Euclidean distances. For a given CDR3 sequence of length *l*, we can then construct its amino acid composition matrix **C** as follows: [*f* (*aa*_1_) *f* (*aa*_2_) *· · · f* (*aa*_*l*_)]^*T*^, resulting in an *l* by *n*_comp_ matrix. **C**. The final embedding is computed as vec(**P***·***C**), providing a vector that combines amino acid compositional information with their positions in the sequence, and is independent of TCR length *l*.

**Figure S2:**
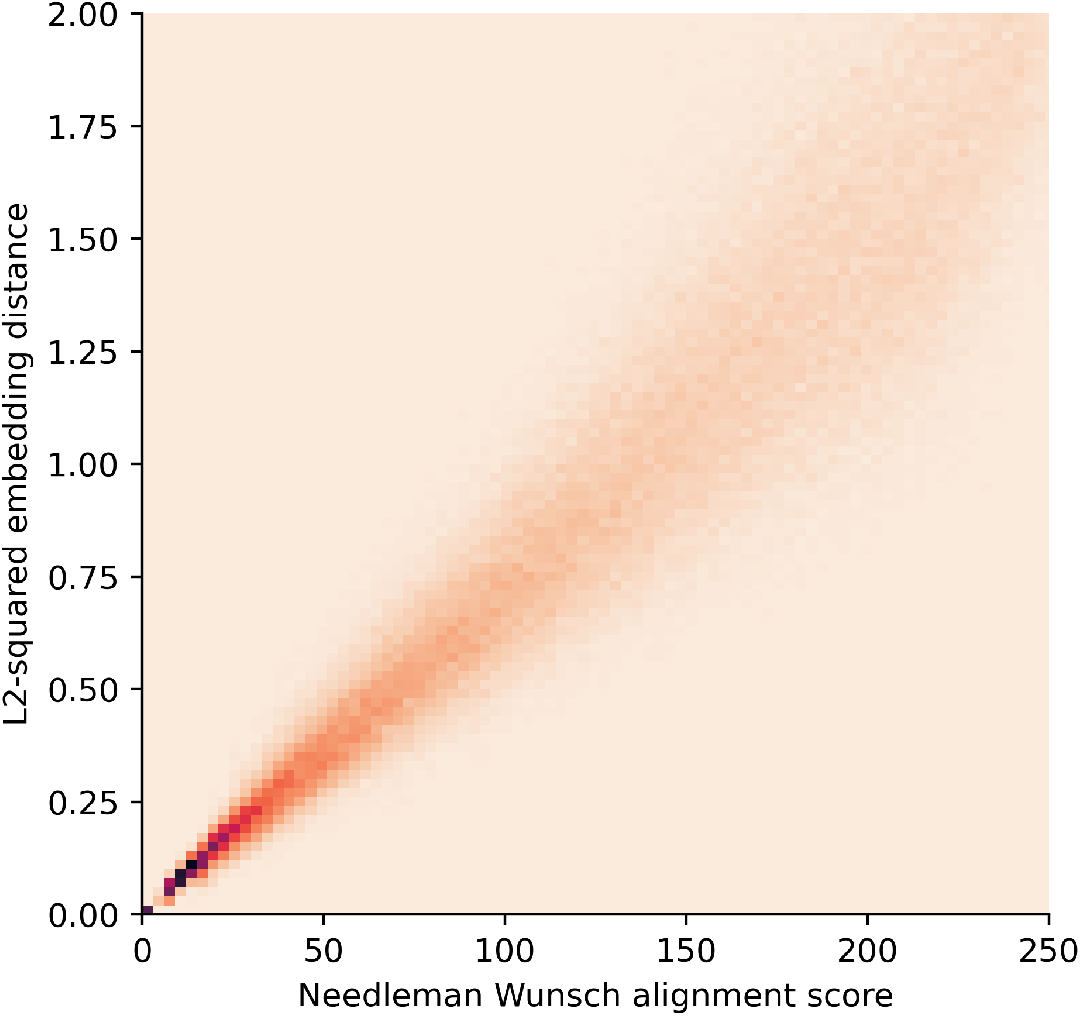
The proximity of a TCR pair’s embeddings closely mirrors their alignment score. We assessed how the distance betweeen TCR sequence embeddings relates to their actual distance. As a ground truth, we measured TCR distance by reciprocal Needleman-Wunsch alignment and BLOSUM62-scoring: dist(*a, b*) = NW(*a, a*) + NW(*b, b*) *−* 2 *\** NW(*a, b*). The NW scores were computed using the with the Needleman-Wunsch alignment and BLOSUM62 scoring from the parasail library (Daily, 2016), using a gap penalty of 4 and an extend penalty of 2. To create a uniform test dataset spanning the entire spectrum of TCR similarity, we randomly selected sequence pairs from the VDJdb and IEDB databases. Specifically, we ensured that each discrete NW-score *≤* 250 was represented exactly 250 times. TCR sequences were embedded using the default parameters, and the embedding distance of a sequence pair was computed as their squared Euclidean (L2-squared) distance. Remarkably, there exists a strong correlation between embedding and alignment distance, as evidenced by a Pearson correlation coefficient *r* of 0.955. This correlation is particularly pronounced for highly similar TCRs, enabling RapTCR’s accurate and efficient vector-based nearest neighbor searches.

**Figure S3:**
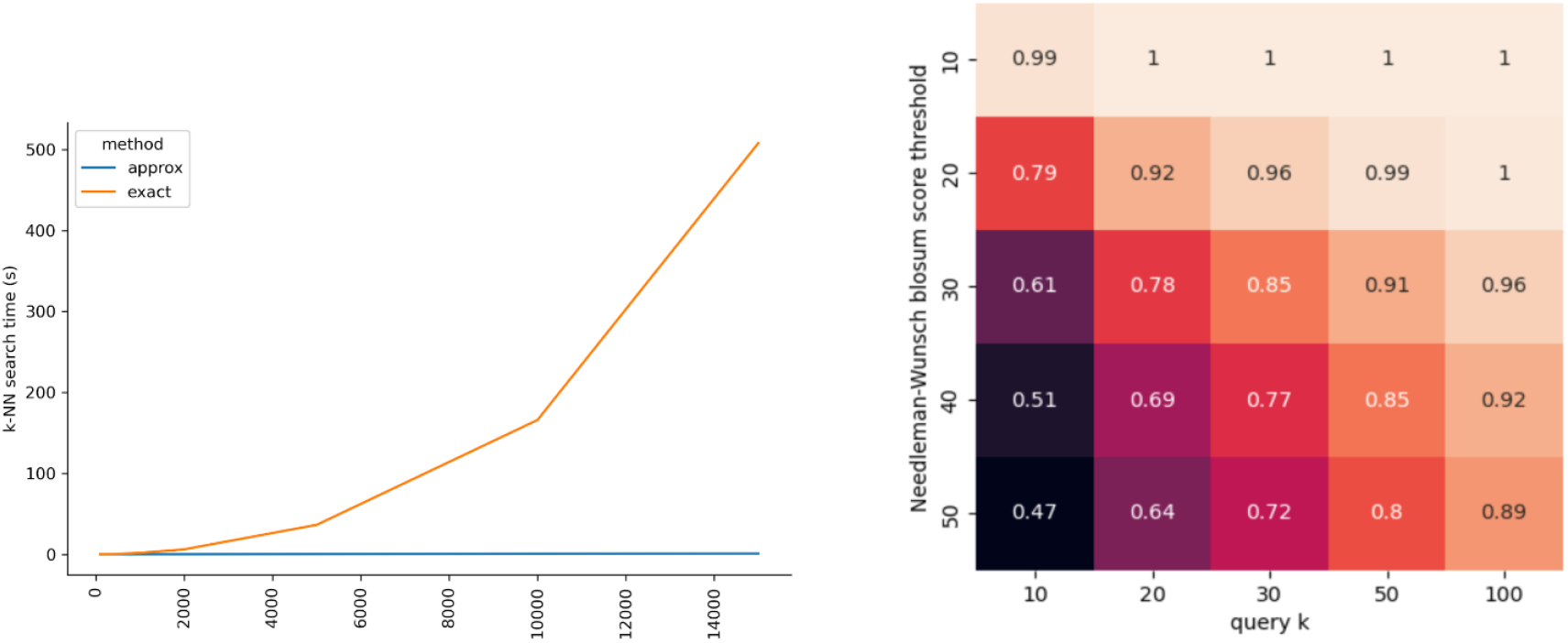
RapTCR finds highly similar sequences with high recall at linear speed. A. Comparative analysis of the speed of k-nearest neighbour (*k*-nn) graph construction with increasing repertoire size (*k*=100). The orange line represents the time needed for an exact search, i.e. brute-force computation of the NW-distance for each sequence pair, and selecting the top-*k* closest pairs. The blue line represents the speed for *k*-NN when using the default exact Faiss-based search implemented in RapTCR. For both methods, parallelism was used with 8 threads on a laptop CPU (Intel i7-12800H). **B**. Recall analysis. As a ground truth, true positives are defined as the 10 TCR pairs with lowest NW-score, if their NW-score *≤* the set score threshold (y-axis). The recall@k for these true positives was determined for the top *k* retreived approximate matches.

**Figure S4:**
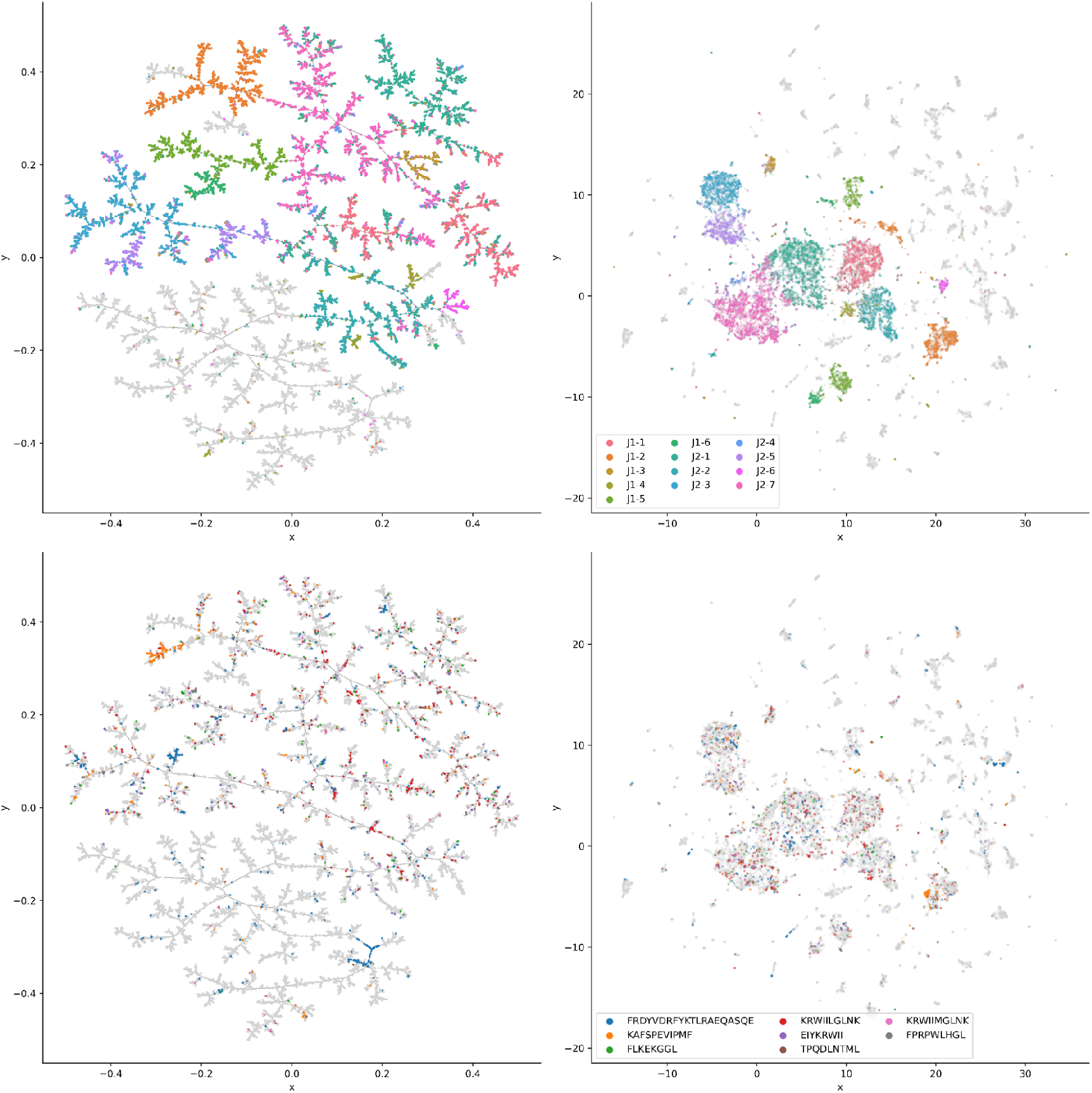
RapTCR visualizations retain global and local similarity features. In this example, the VDJdb dataset was plotted both using the MST-based (left) and UMAP layout (right). **A**. Major global features preserved in both visualizations are the TCR chain type (TRA or TRB), and the J-gene. **B**. Arguably the most significant feature of a TCR sequence is its epitope-reactivity. Hence, TCR reactivity for the eight most common HIV-1 epitopes was overlaid as a color layer on the RapTCR plots. Notably, most of these HIV-1-annotated TCRs are CDR3 beta chains, except a large FRDYV..-reactive TCR group. Interestingly, for most epitopes, a large fraction of their annotated TCRs cluster densely together, indicating the presence of a group of highly similar receptor sequences that recognize this epitope. In contrast, KRWILL-reactive TCRs are more sparsely spread out over the sequence space in smaller groups, indicating that they can be recognized by a more diverse set of CDR3 sequences.

